# Ultrastructural Organization of the Honeybee Blood-Brain Barrier and Comparison with Age

**DOI:** 10.1101/2024.09.27.615080

**Authors:** Tyler Quigley, Gro Amdam

## Abstract

The blood-brain barrier is an essential feature of the most animal nervous systems, adapted to maintain brain homeostasis across species occupying unique and dynamic environments. Understanding the structural and functional diversity of animal blood-brain barriers can aid our understanding of how this structure contributes to important areas of neuroscience such as behavior, health, and disease. The honeybee worker is a well-established model for exploring these dimensions of brain function, however, the honeybee worker blood-brain barrier has yet to be described. Here, we present the first global anatomic analysis and description of the honeybee worker blood-brain barrier and compare key ultrastructural features between two age groups. We describe the cellular makeup of the honeybee worker blood-brain barrier, patterns of heterogeneity in barrier structure throughout the brain, and highlight important areas of further study.

## Introduction

Behavior arises from intricate electrical and chemical communication networks among neurons in the central nervous system (CNS). The distinct electrochemical physiology of neurons demands both a constant supply of energy and a strict ionic environment. To meet these demands, nearly all animal taxa have developed a cellular barrier that compartmentalizes neuronal cells from vascular spaces (Abbott, 1992; Abbott *et al*., 2010). This barrier is most often, including in this article hereafter, referred to as the ‘blood-brain barrier’, including in reference to animals which have hemolymph rather than blood. The primary functions of the blood-brain barrier are largely consistent across species; it acts as a physical and chemical barrier to molecules in circulation to the CNS, while selectively transporting in vital biomolecules through a network of transporters (Weiler *et al*., 2017; Dunton *et al*., 2021).

The earliest conception of the blood-brain barrier was built based on series of experiments by research groups who examined the fate of dyes and other agents injected into the blood, brain, and spaces around the brain in animal models (Ehrlich, 1885; Lewandowsky, 1900; Goldmann, 1913; Stern and Gautier, 1918). These early 20^th^ century experiments demonstrated that molecules could not necessarily flow freely between the brain and periphery, suggesting the existence of a selective barrier around the brain (Stern, 1934; Vein, 2008; Pivoriūnas and Verkhratsky, 2021). Subsequent advances in electron microscopy led to the localization of the blood-brain barrier to the blood vessels in the brain, the modified endothelial cell composition of this barrier, and the presence of tight junctions in the intercellular space. The primary goal of blood-brain barrier research throughout the 20^th^ century subsequently became to understand its role in neurological health and disease, and how it can be manipulated to improve drug delivery (Hurst and Davies, 1950; Saunders *et al*., 2014; Dunton *et al*., 2021). The model organisms most often employed in this research were chosen for their translational potential, thus composed primarily of mammals such as rodents, rabbits, pigs, dogs, and humans (Ribatti *et al*., 2006).

Despite the progress made in understanding the significance of the blood-brain barrier to human health, the bias towards mammalian models has led to a limited understanding of the blood-brain barrier as an essential feature of the nervous system across the animal kingdom. Indeed, blood-brain barriers are nearly ubiquitous across animals, and the modern phylogenetic view of blood-brain barrier evolution is that it evolved independently at least three times (Abbott, Lane and Bundgaard, 1986; Bundgaard and Abbott, 2008). It has been hypothesized that increasing complexity in the central nervous systems selects for an increasingly complex diffusion barrier around the brain (Dunton *et al*., 2021). However, the cell types capable of forming such a barrier in the face of this selection pressure have varied across taxa, which has led to the evolution of blood-brain barriers with convergent functions yet diverse structures across lineages. Thus, despite these morphological differences between taxa, non-mammals can provide fundamental insights into blood-brain barrier physiology translatable to mammals and even humans (DeSalvo *et al*., 2014; Weiler *et al*., 2017). In particular, studies of the insect blood-brain barrier studies have revealed fundamental aspects of blood-brain barrier structure, transport mechanics of endogenous and exogenous molecules between the nervous and circulatory systems, and the role of the blood-brain barrier in neurodegenerative diseases (Carlson *et al*., 2000; Andersson *et al*., 2013; Contreras and Klämbt, 2023).

The insect blood-brain barrier is composed of an acellular fibrous structure called the neural lamella, and two subtypes of surface glia, perineurial glia (PG) and subperineurial glia (SPG). The neural lamella is a fibrous matrix devoid of cells that envelops both the central and peripheral nervous system. This electron-dense layer is believed to consist of extracellular matrix components, such as proteoglycans, mucopolysaccharides, and collagens, providing essential structural support to the neural tissue (Dunton *et al*., 2021). PG are positioned directly beneath the neural lamella forming the outermost cellular layer of the brain. They assume spindle-like shapes and extend slender protrusions towards neighboring PG cells and the underlying SPG layer. The precise functions of PG are not well understood, though it is possible that they act as both an initial diffusion barrier and a hemolymph nutrient sensor (Contreras and Sierralta, 2022). SPG are large, flat, polyploid cells that form extensive interdigitations with neighboring SPG, which are further connected by pleated septate junctions. SPG thus form the tight diffusion barrier around the brain and are the likely location of active transport systems that shuttle molecules between the brain and blood. Collectively, these blood-brain barrier components preserve brain homeostasis in similar ways as the mammalian blood-brain barrier; though the cell types and morphology differ, chemoprotective and small molecule control pathways are largely conserved among the taxa (DeSalvo *et al*., 2014).

Outstanding questions about blood-brain barrier physiology that are particularly suited for study in insects include how heterogeneity in blood-brain barrier composition is organized, how this heterogeneity is used to maintain brain homeostasis, and how heterogeneity changes over the lifespan of a species (Noumbissi, Galasso and Stins, 2018; Chen *et al*., 2019; Griffith *et al*., 2020). The insect brain is small and thus amenable to global analyses of blood-brain barrier heterogeneity. The insect blood-brain barrier itself is easily localized, as it exists as an outer sheath of the brain, and blood-brain barrier components are easily identified in both light and electron microscopy. The *Drosophila* system offers an expansive genetic toolkit for probing blood-brain barrier physiology, however, a multispecies approach to questions of barrier heterogeneity can reveal common and divergent patterns across species that occupy different niches and with different life histories.

Honeybee (*Apis mellifera*) workers are a well-established neurobiological model frequently used to explore fundamental nervous system properties due to their wide repertoire of advanced behaviors, well-described neuroanatomy, and array of experimental tools and paradigms (Scheiner *et al*., 2013; Quigley, Amdam and Rueppell, 2018; Groh and Rössler, 2020; Habenstein *et al*., 2023; Kraft *et al*., 2023). Honeybee workers are eusocial, nonreproductive members of societies composed of up to 50,000 members, including a single queen and a few thousand males (Winston, 1987). Over a century of research on honeybee worker behavior has revealed that this insect caste can express advanced learning, memory, communication, time tracking, navigational abilities, among other behaviors that drive their collective handling of essential colony functions (Seeley, 1986; Menzel, 2012; Avarguès-Weber and Giurfa, 2013; Scheiner *et al*., 2013; Cook *et al*., 2020; Siefert, Buling and Grünewald, 2021). Simultaneously, extensive neuroanatomical analyses of the worker brain have characterized the neuronal architecture of the worker brain and has resulted in the construction of a detailed brain atlas (Kenyon, 1896; Rybak, 2012; Habenstein *et al*., 2023). These cumulative efforts have succeeded in connecting aspects of the advanced cognitive and behavioral abilities of honeybee workers to discrete regions, neuropils, and neuronal networks in their brain (Menzel, 2001; Menzel and Giurfa, 2001; Sommerlandt *et al*., 2019; Paoli and Galizia, 2021). Neurons typically compose 90% of the cells in the insect brain, while the other 10% are composed of glial cells, of which the insect blood-brain barrier is composed (Pogodalla, Winkler and Klämbt, 2022). Detailed investigations of glial spatial patterns throughout the brain may lead to a more comprehensive model of the honeybee worker brain and encourage future studies to better elucidate their functions.

Here, we present a global anatomical characterization of the ultrastructure of the worker honeybee blood-brain barrier. We employed a systematic uniform sampling technique paired with transmission electron microscopy (TEM) to unbiasedly sample brain ultrastructure. We focused our analysis on the most outer layer of cells where the insect blood-brain barrier is located (Figure 1) (Schofield and Treherne, 1984; Desalvo *et al*., 2011). To gain further insights into how aging may influence the structure of the honeybee blood-brain barrier, we conducted a comparative stereological analysis of mitochondrial density, nuclear density, and barrier depth in young and old foragers. Although both age demographics constitute the final stages of the honeybee lifespan, senescent patterns in the honeybee brain typically manifest in old foragers, particularly those that have been foraging for 14 or more days (Seehuus, Krekling and Amdam, 2006; Behrends *et al*., 2007; Wolschin, Münch and Amdam, 2009; Munch *et al*., 2013). By comparing these two groups, we aimed to uncover structural changes in the blood-brain barrier that may occur during the aging process.

**Figure 1.**
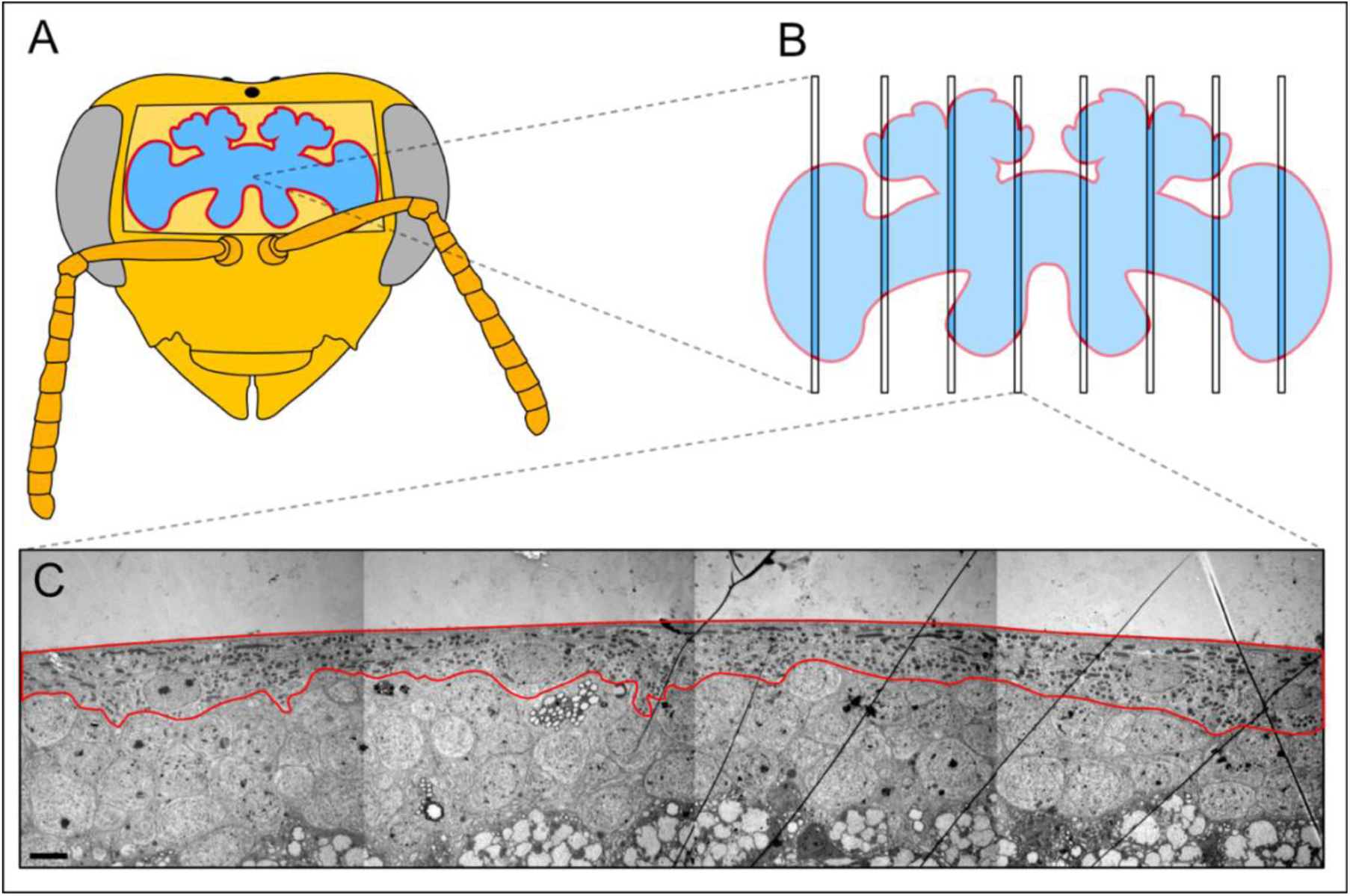
(A) Illustration of a honeybee head with a window cut out to reveal the brain. (B) Honeybee brains were dissected, prepared for electron microscopic imaging, and then systematically uniformly sampled so that every point in the tissue has an equal chance to be sampled. (C) Each section obtained from the brain was imaged in overlapping portions and stitched together to create a montage of the entire blood-brain barrier as sectioned at that point in the brain. The red outline in (A) & (B) represents the blood-brain barrier as the most outer layer of the honeybee brain. The red outline in (C) surrounds the blood-brain barrier, which includes the neural lamella, perineurial glia, and subperineurial glia (scale bar: 2 μm).

## Methods

### Animals

Honeybee workers were obtained from two host colonies from the apiary at Arizona State University Polytechnic Campus in the spring of 2017. Approximately 1,500 foragers were marked with a paint pen upon their return from foraging. Fourteen days later, another subset of 200 foragers were marked upon their return from foraging with a paint pen of a different color. To avoid accidentally marking non-foragers, all forager marking was completed during active foraging hours in the early morning before orientation flights begin. Returning foragers typically fly in a straight line for the colony entrance and are weighed down either by a full crop or pollen load. This results in recognizable behavioral and flight patterns which were used to determine which honeybees at the entrance are foragers, and to minimize the marking of non-forager honeybees. The following day (15 days since first marking), five bees of both groups were collected. The bees collected with the first color mark had foraged for at least 15 days, here considered old foragers. The bees collected with the second color mark had foraged for at least one day, here considered young foragers. Although it is possible that individuals that had been foraging for <15 days were captured in this group, the chances are minimal. After ten days of foraging, the mortality rate of foragers increases dramatically with each subsequent day (Dukas, 2008). At 15 days foraging, <10% of foragers can typically be recovered, reflecting their low representation in the population (Wolschin, Münch and Amdam, 2009). This fact paired with the small sample size needed for this study nearly guarantees that the young forager specimens in this study are <15 days foraging. Three foragers from each group were included in the present study, however, one young forager sample was eliminated prior to analysis due to poor tissue preservation.

### Electron Microscopy

All sample preparation, sectioning, and imaging took place in the ASU Eyring Materials Center Life Sciences Electron Microscopy Lab. Honeybee brains were dissected in a fixative of 2% PFA/2.5% glutaraldehyde in Sorenson’s buffer (pH 7.2), and stored in this fixative for 24 hours at 4°C. After washing in Sorenson’s buffer (3X 10 minutes), brains were post-fixed in 1% OsO4 for 1 hour at room temperature. After another wash in Sorenson’s buffer (2X 10 minutes) and diH_2_O (3X 10 minutes), brains were en bloc stained with 2% uranyl acetate for 1 hour. After a final wash in diH_2_O (3 x 10 minutes), brains were dehydrated in a graded series of ethanol washes (20%, 40%, 60%, 80%, 95%, 3X 100%; 10 minutes each). Brains were then infiltrated with an ascending series of Spurr’s resin mixed with ethanol (1:3, 2:2, 3:1, 3X 4:0; 4+ hours each) on a rotating wheel. Each brain was then placed in a random orientation into the lid of Beem capsule filled with Spurr’s Resin, and polymerized for 30 hours at 60°C.

Once hardened, the circular block was spun by hand on a flat surface (e.g. tabletop) to randomize the orientation of the tissue with respect to a pre-determined edge of said surface, and a picture was taken of the orientation. The circular block was trimmed down to the tissue and re-embedded in a flat mold in the randomized orientation. The final blocks were manually trimmed with a razor blade to a block face which included the edge of the brain, where the blood-brain barrier is located. Two sets of 5 ultrathin sections (75 nm) were taken from these blocks every 1 mm along the blood-brain barrier along the randomly oriented axis of the brain. Sections were collected onto formvar-coated copper slot grids. Grids were post-stained for 12 minutes in 2% uranyl acetate (Electron Microscopy Sciences) in 50% EtOH and 9 minutes in Sato’s lead citrate (Electron Microscopy Sciences).

### Stereological Imaging with TEM & Image Stitching

Tissue sections were imaged using a Phillips CM12 TEM at a magnification of 4400X. The resulting micrographs were approximately 21 μm^2^. Consecutive micrographs were taken such that each micrograph overlapped slightly with the previous micrograph. This enabled the micrographs to be stitched together to create a large montage spanning the entire length of the blood-brain barrier captured in each section (see Figure 2A and 2.4A-2.4E). Images were overlayed and stitched together using Affinity Photo (Version 1.10.5; Serif, 2022).

**Figure 2.**
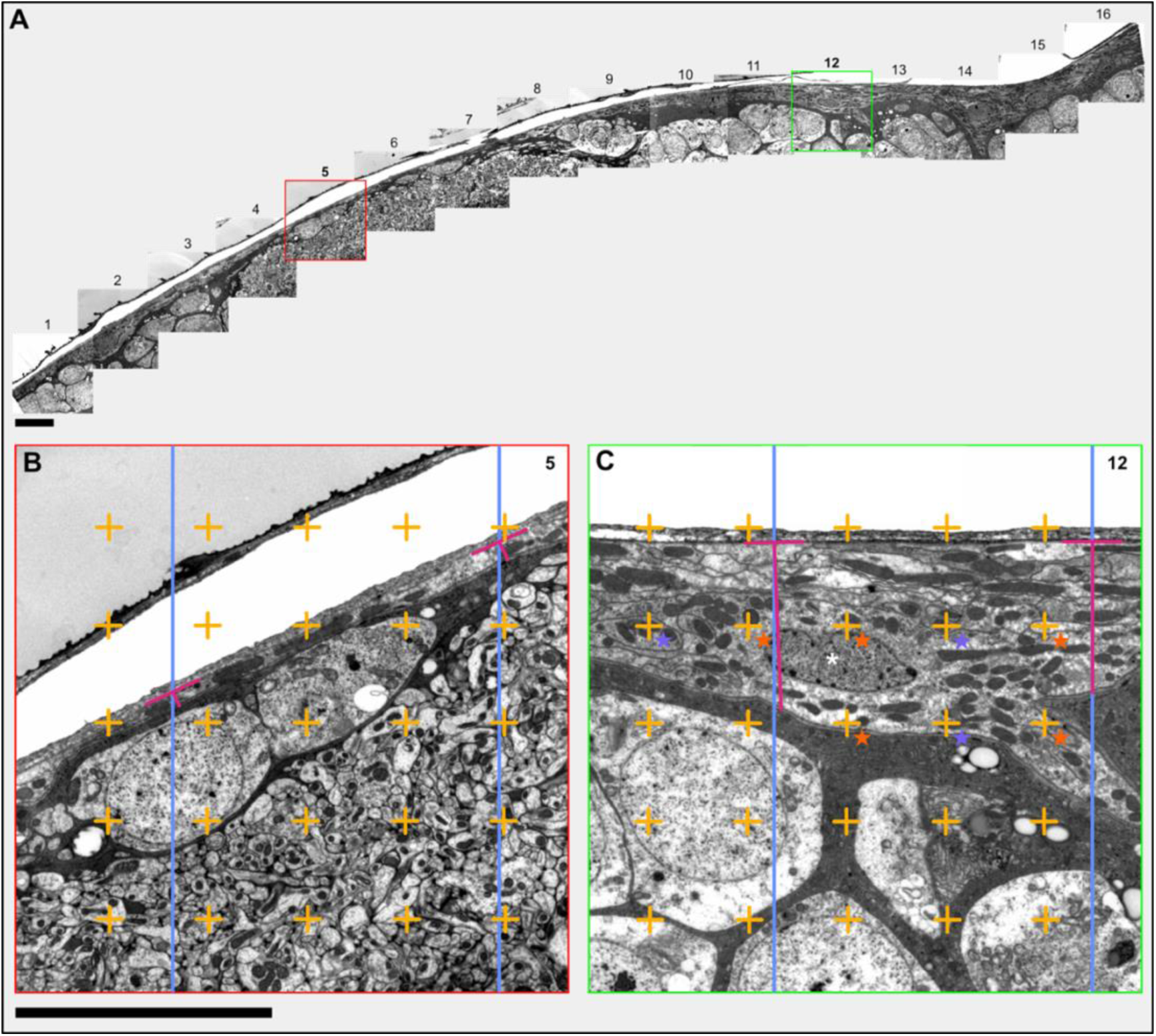
Example of a montage and illustration of stereological analysis tools. (A) Montage of 16 micrographs captured of the blood-brain barrier and underlying layers of cells. The blood-brain barrier is the most outer layer of cells; compare to the encircled cells in Figure 1C. Glial cells are more electron dense than neurons, thus show up darker in transmission electron micrographs. Note the diversity in cellular composition below the blood-brain barrier. (B) An enlarged view of micrograph 5 from (A), depicting a thin stretch of blood-brain barrier composed of a single SPG cell, with a depth of approximately 0.5-0.9 μm. (C) An enlarged view of micrograph 12 from (A), depicting a thicker stretch of blood-brain barrier with a depth of approximately 5.8-6.5 μm, and containing a nucleus (white asterisk). (B, C) To measure mitochondrial and nuclear density, a grid (light orange crosses) was overlayed at random over the entire montage. A count was made wherever the bottom right quadrant corner of a cross intersected a mitochondrion (purple star), nuclei, or the reference space (orange star). No intersections occurred in (B), and no intersections with a nucleus occurred in either section. To measure barrier depth, a uniformly spaced set of lines (blue vertical lines) was overlayed at random over the entire montage. At the point where the line intersected the apical side of the blood-brain barrier, a T figure (magenta) was made. The horizontal line of the T was drawn parallel to the apical border, and the vertical line was drawn to the basal border. The vertical line was measured as barrier depth. Scale bars: 10 μm.

### Tissue preservation optimization

We made a notable observation regarding the preservation of blood-brain barrier ultrastructure during tissue processing for TEM. Tissue sections where the neural lamella was disrupted also exhibited a disrupted blood-brain barrier. Consequently, such sections were excluded from the analysis, as the distorted blood-brain barrier structures observed were often shrunken, dissolved, or otherwise compromised, failing to accurately represent the true blood-brain barrier architecture. To enhance the preservation of the blood-brain barrier structure across samples, we implemented modifications in the experimental procedure. This included introducing the fixative to the brain at an earlier stage during the dissection process and exercising extra care in the physical handling of brain tissue. Specifically, after decapitation of the head and removal of antennas, the head was placed into fixative for approximately 30 minutes before the brain was dissected from it within a ∼2 mL droplet of fixative. Examples of extra handling precautions taken include removing as much extraneous tissue from around the brain while it is still within the head capsule, exclusively grasping the retina with forceps while handling the brain and scooping the brain with forceps rather than pinching whenever possible. These minor fixation and handling adjustments yielded greatly improved results in maintaining the integrity of the blood-brain barrier structure throughout the analysis.

### Stereological Analysis in ImageJ

The stereological analyses below were performed using the Fiji distribution of ImageJ and the multipurpose grid macro version 1.0 (A. Mironov, University of Manchester; https://imagej.nih.gov/ij/macros/Multipurpose_grid.txt) (Mironov, 2017). A constant scale (area per point) of 48.1251 pixels/um was used.

#### Mitochondrial (V_vmit_) & Nuclear (V_vnuc_)Volume Fraction

The technique used here is derived from Delesse’s geological principle which states that the area fraction of a mineral is equal to its 3D volume fraction (Mayhew and Orive, 1974). This principle can be applied to any compartmentalized 3D volume, such as biological tissue, and is used for many such applications (Hartung, Kirkton and Harrison, 2004; Marcos, Monteiro and Rocha, 2012; Santuy *et al*., 2018). The multipurpose grid plugin was used to overlay a grid in random orientation over the images (Figure 2B and 2.2C). The cell counter tool was used to count points that landed on mitochondria, nuclei, and the reference space across the entire image. Reference space was defined as the blood-brain barrier cell layer, defined here as the space between the apical membrane of PG and the basal membrane of the SPG.

#### Barrier Depth

The multipurpose grid plugin was used to overlay a random set of lines perpendicular to the general direction of the blood-brain barrier in the montaged micrographs (Figure 2B & 2C). At each line intersection, a T-figure was drawn; the horizontal line of the T was drawn parallel to the neural lamella, and the perpendicular vertical line drawn between this line and the basal border of the SPG layer. The length of the perpendicular vertical line was measured as barrier depth. Between 51-76 depth measurements were made per individual, resulting in a total of 127 barrier depth measurements among young foragers and 167 barrier depth measurements among old foragers.

### Statistical Analysis

Stereological probe counts were used to compare BBB structural parameters between young and old foragers. Comparisons of V_vmit_, V_vnuc_, and barrier depth between age groups were made using t-tests. The barrier depth dataset was skewed to the left, reflective of large variation in the blood-brain barrier depth. This dataset was normalized using a log-transformation and a Welch’s t-test to account for unequal variances between the two comparison groups. The log-transformed data is presented as boxplots depicting the median (bold center line), interquartile ranges (span of box), range within 1.5 times the interquartile range (whiskers) and outliers. All statistical analyses were performed in R.

## Results

### Sampling

The SURS sampling and image stitching technique yielded a total of 17 reconstructed images, each composed of between 8 and 37 overlapping micrographs, and a median micrograph count of 18. Per individual 3 - 4 montages contained sufficiently preserved tissue for stereological analysis.

### Neuroanatomy of the honeybee blood-brain barrier

As in other insects, the honeybee blood-brain barrier is composed of a neural lamella, PG, and SPG (Figure 3). Throughout our extensive imaging we captured a variety of cell types and substructures which border the basal surface of the honeybee blood-brain barrier. We identified cell types and subcellular cell structures based on their location and morphology as reported in the extensive literature on insect central nervous system structure (Ito *et al*., 2014; Freeman, 2015; Kremer *et al*., 2017; Kaiser *et al*., 2022; Habenstein *et al*., 2023).

**Figure 3.**
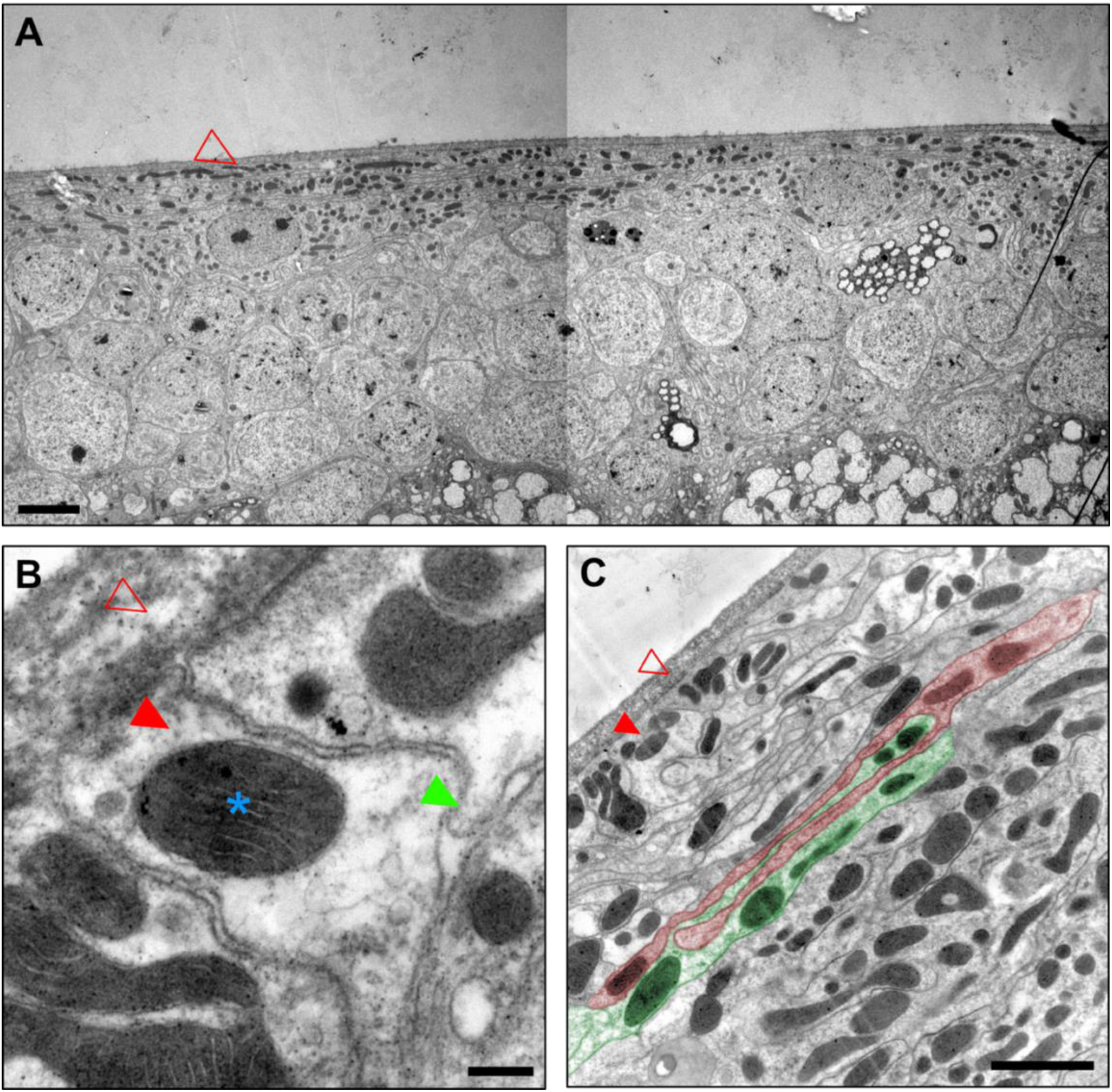
The primary components of the honeybee blood-brain barrier. (A) An overview image of the blood-brain barrier reveals its smooth apical edge facing the hemolymph and covered by the neural lamella (red outlined arrow all panes) and convoluted basal edge interfacing with the underlying cortex glia and neuron layer. The blood-brain barrier is mitochondrially dense (dark, bulbous shapes, denoted in (B) with blue asterisk), likely reflecting a high level of active transport between the hemolymph and brain. (B) A higher magnification image of a perineurial glial cell (solid red arrow in B and C). The intercellular space between perineurial glia runs perpendicular to the neural lamella, which may allow passage of small molecules to the subperineurial glia layer. The basal edge of perineurial glia overlap with each other (green arrow) and the underlying subperineurial glia. (C) Subperineurial glia compose the bulk of the blood-brain barrier and form the physical diffusion layer that prevents the free passage of molecules in the hemolymph from entering the brain. Here, two subperineurial glial cells are highlighted in red and green to accentuate the extensive interdigitation between neighboring SPG cells. (A and C) scale bar 2 μm. (B) scale bar: 0.2 μm.

### Neural lamella

The neural lamella was identified throughout the honeybee brain, and its preservation as the outer layer of fixed tissue used to determine the overall quality of tissue preservation. Under the TEM, it appears as a blurry, mesh-like network of electron-dense fibers tightly associated with the underlying cells. The only other structure we identified within the neural lamella are tracheoles (Figure 4D & 4E). We observed that the thickness of the neural lamella varies between 0.5 μm and 3 μm, with greater thickness observed at points where the tracheal network enters the brain (Figure 4E).

**Figure 4.**
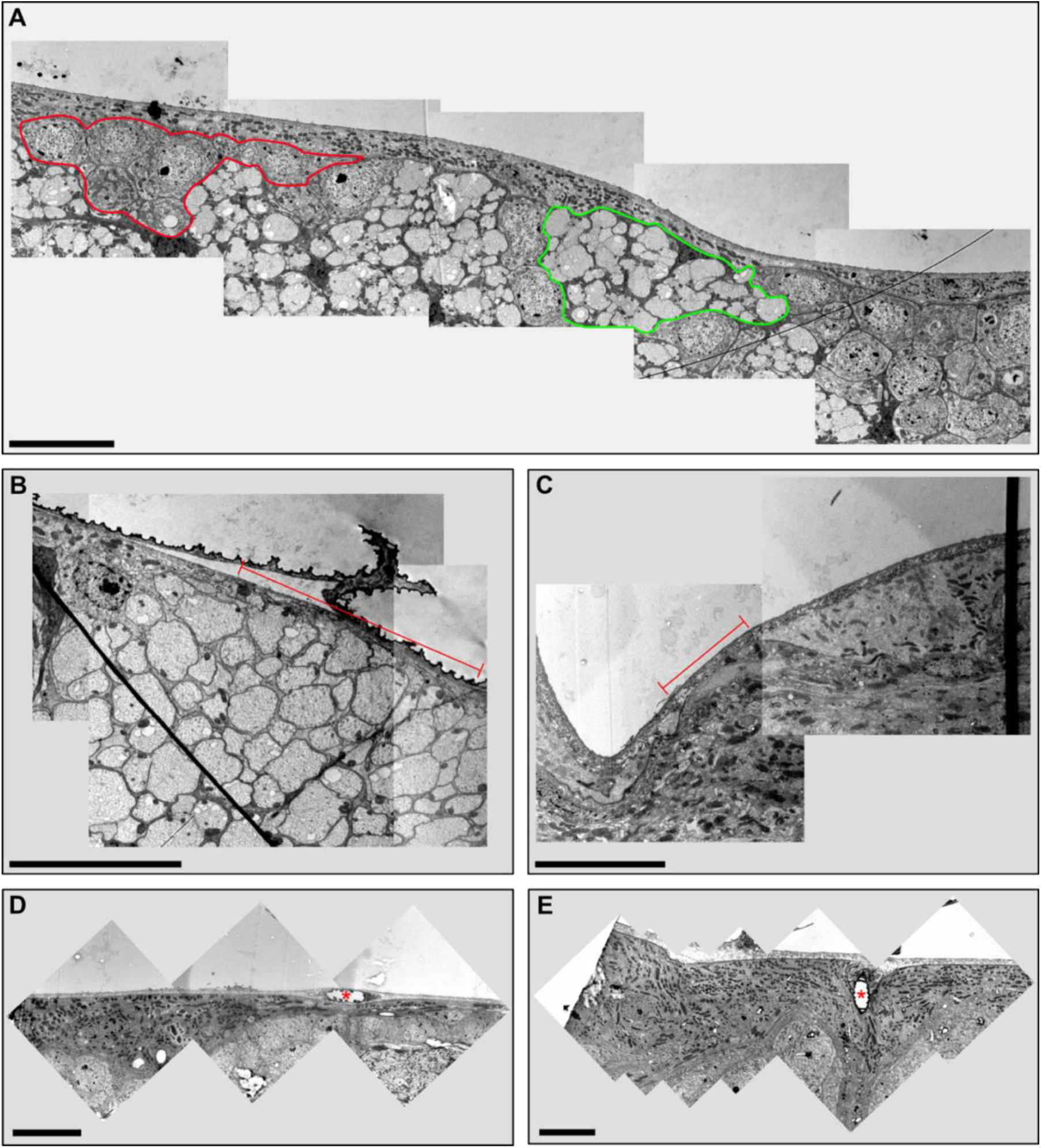
Various points of note in the honeybee blood-brain barrier. (A) A montage depicting variation in glial cell types and neuronal structures bordering the basal edge of the honeybee blood-brain barrier. Cortex neuronal cell bodies surrounded by cortex glia are encircled in red. Dendritic or axonal arbors surrounded by ensheathing glia are encircled in green. (B) Micrographs depicting a region of blood-brain barrier consisting of a single surface glial layer between the neural lamella and dendritic or axonal arbors (red line). (C) Micrographs depicting a region where there is a lack of surface glial cells between the neural lamella and non-blood-brain barrier cell types (red line). (D, E) Montages depicting tracheoles intersecting with the blood-brain barrier. Note the depth and mitochondrial density of the blood-brain barrier surrounding the tracheole entering the brain in (E). Scale bars: 10 μm.

### Perineurial glia

PG are one of the two surface glia subtypes that constitute the insect blood-brain barrier. We captured PG in various orientations relative to the section cuts and found that their distribution and morphology in the honeybee brain align with the descriptions provided in the existing literature on insect PG (Awasaki *et al*., 2008; DeSalvo *et al*., 2014; Freeman, 2015). The apical surfaces of PG cells appear flat, running parallel to the basal border of the neural lamella. The intercellular space between adjacent PG cells open towards the neural lamella. Along their longitudinal axis, PG cells form overlapping ledges with neighboring cells while extending elongated, slender protrusions towards both neighboring PG cells and the SPG layer (Figure 3B). Our observations align with the known characteristics of PG in insect blood-brain barrier architecture.

### Subperineurial glia

SPG are the second surface glia subtype that comprises the insect blood-brain barrier. Our analysis of the honeybee blood-brain barrier confirms that SPG form a continuous and highly interconnected cellular layer beneath the PG layer. SPG cells are characterized by their flattened and elongated shape, and their cell membranes exhibit extensive folding and interdigitation with neighboring SPG cells (Figure 3C). This interdigitation creates a complex network of tight junctions, resembling a maze, which serves as a physical barrier, preventing the passage of molecules from the hemolymph into the brain. At the ultrastructural level, SPG cells appear flattened and possess numerous microvilli. They are densely packed with mitochondria and often contain one or more nuclei. Through extensive sampling, we have observed the ubiquitous presence of SPG cells around the brain, constituting the majority of the blood-brain barrier depth, ranging from less than 0.5 μm to 10 μm in thickness.

### Single surface glial layer

While both PG and SPG are found throughout the brain, our imaging reveals the presence of blood-brain barrier locations where only a thin, single layer of protrusion from a surface glial cell separates the neural lamella from the underlying CNS cells (Figure 2B & 4B). We also identified at least one point in the blood-brain barrier where surface glia are entirely absent, and another cell type interfaces directly with the neural lamella (Figure 4C).

### Associated cell types

Our random sampling technique was designed to grant every point on the brain surface near-equal probability of being sampled. This enabled us to observe the relationship of the honeybee blood-brain barrier to the diversity of cell types located immediately deep to the barrier. The composition of cells underlying the blood-brain barrier are extremely diverse, however, we observed no instances where the blood-brain barrier was in direct contact with a neuron. As a rule, the basal border of the blood-brain barrier always interfaces with another glial subtype. Identification of glial cell types on ultrastructure alone is difficult, however, the morphology and neuronal association of the glial cells that border the blood-brain barrier fit the profile of cortex glia and ensheathing glia (Kremer *et al*., 2017; Pogodalla, Winkler and Klämbt, 2022). Cortex glia surround and fill the space between the neuronal cell bodies of cortex neurons that lie towards the outer perimeter of the brain, while ensheathing glia are electron dense and surround dendritic and axonal regions. We found that cortex glia most often interface directly with the blood-brain barrier, however, some of our images suggest ensheathing glia may share this space in some regions of the brain (Figure 4A).

### Stereological comparison of young and old foragers

The global volume fraction of mitochondria and nuclei within the blood-brain barrier of young foragers (<15 days foraging) and old foragers (≥15 days foraging) is shown in Figure 5. Within the blood-brain barrier of young foragers (n=2), we observed a mitochondrial volume fraction of 36.2 ± 0.04%, and a nuclear volume fraction of 10.2 ± 0.05%. Within the blood-brain barrier of old foragers (n=3), observed a mitochondrial volume fraction of 39.6 ± 0.03%, and the nuclear volume fraction of 10.8 ± 0.01%. The average mitochondrial and nuclear volume fraction across all specimens (n=5) was, respectively, 38.3 ± 3.2% and 10.6 ± 2.8%. We found no significant difference in mitochondrial volume fraction (t(3.98) = 1.75, p = 0.16) nor nuclear volume fraction (t(2.25) = 0.9, p = 0.45) between the two age groups.

**Figure 5.**
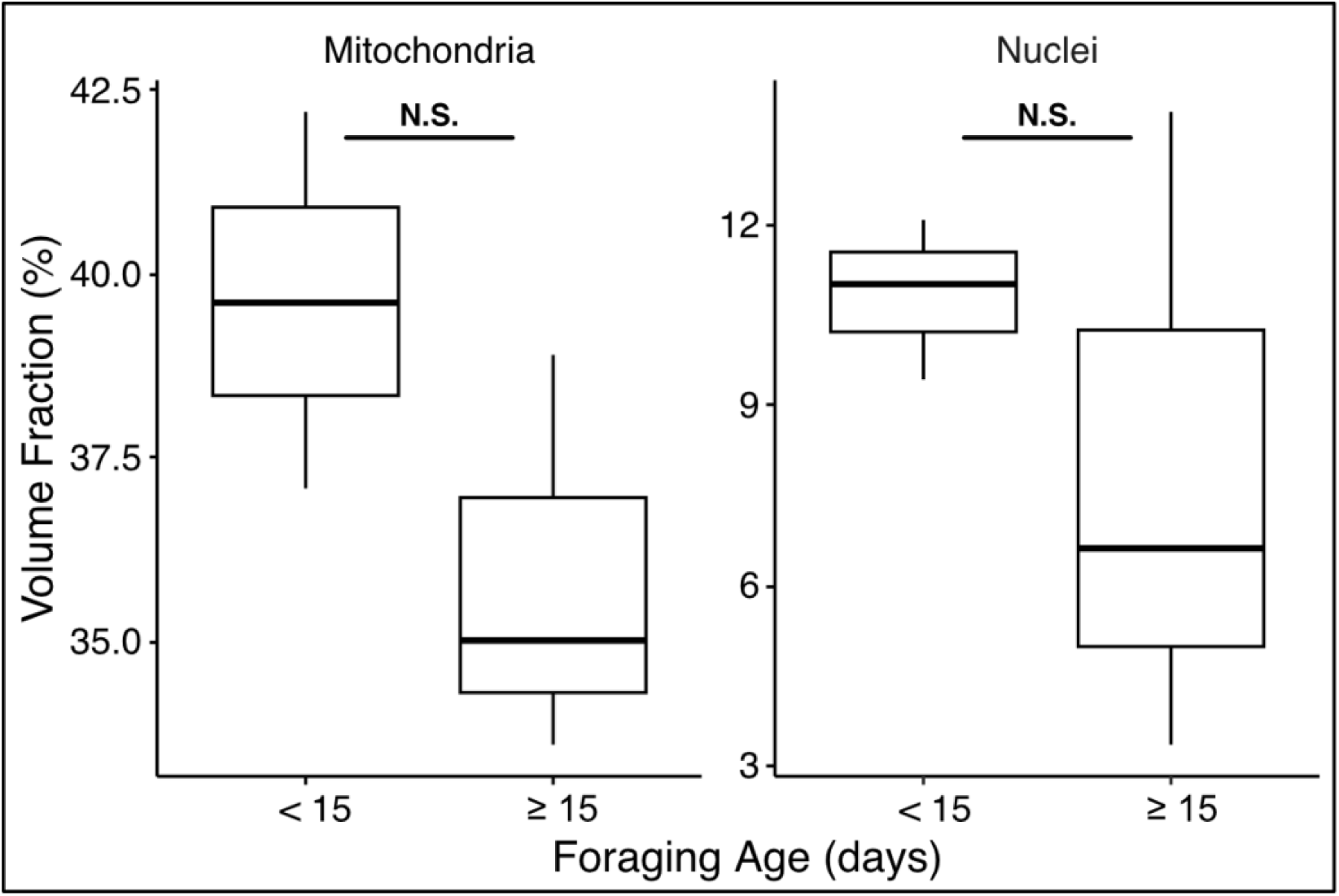
Volume fraction of mitochondria and nuclei in young and old foragers. We found that the mean volume fraction of both mitochondria and nuclei was higher in the younger forager group, however, the differences were not significant. Individual volume fraction measurements were made from 9 montages across 3 young foragers and 9 montages across 3 old foragers. N.S. refers to non-significance. Mitochondria: t(3.9) = 1.75, p = 0.16; Nuclei: t(2.2) = 0.9, p = 0.05.

The depth of the blood-brain barrier varied greatly across all of the specimens (n=5), ranging from a minimum of 0.345 μm to a maximum of 27.8 μm with an average of 5.36 μm. Figure 6 depicts the difference in log-transformed barrier depth (defined as the distance from neural lamella to the border of SPG with the underlying cells) between young and old foragers. Old foragers had a mean barrier depth of 6.32 μm, while young foragers had a mean barrier depth of 4.16 μm. Despite nonsignificant differences between the mitochondria and nuclear volume fraction of these two groups, we observed that the depth of blood-brain barrier in old foragers was significantly larger than that of young foragers (t(285.4) = 3.59, p<0.0005).

**Figure 6.**
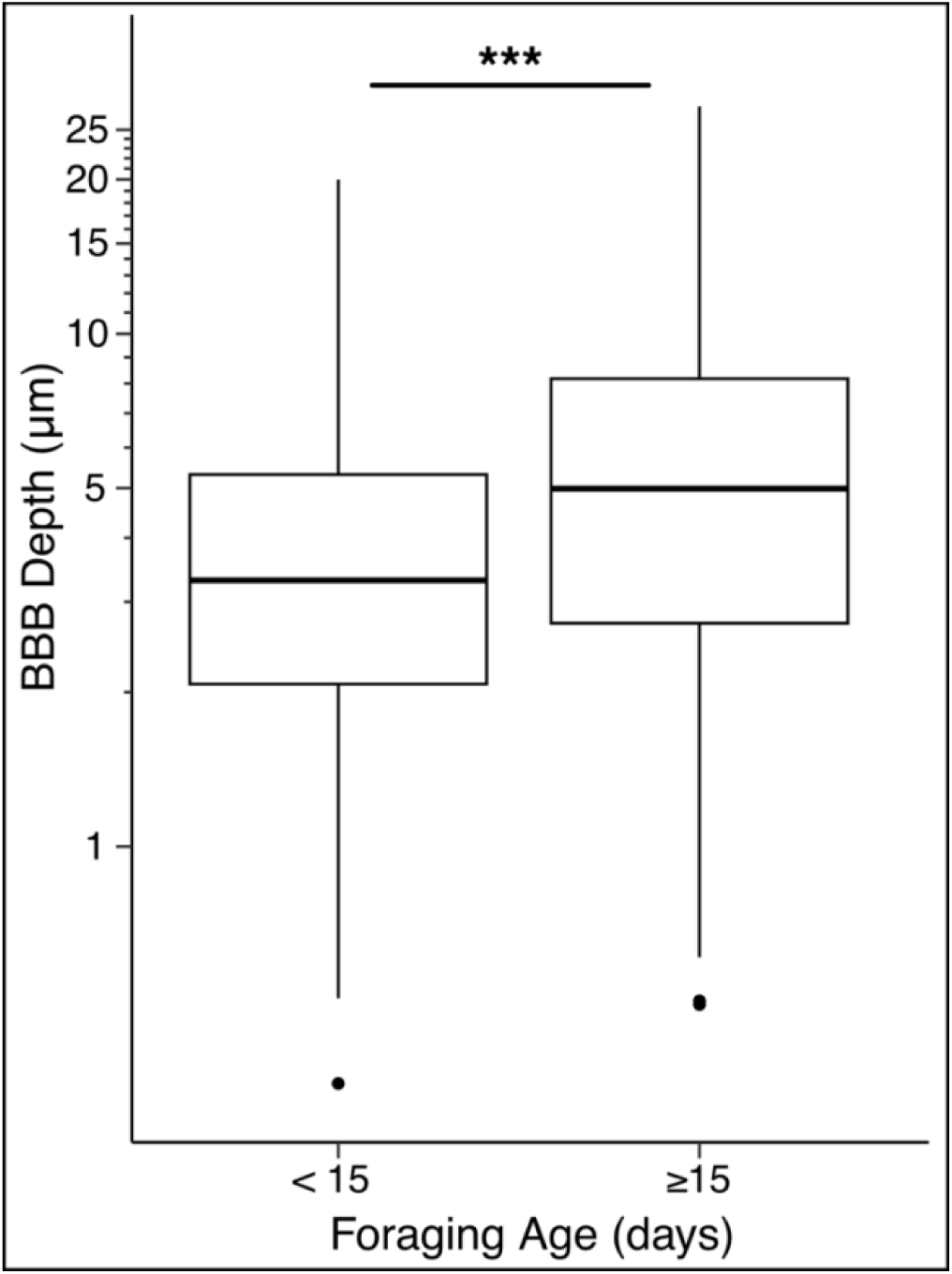
Comparison of barrier depth between young and old foragers. Barrier depth is defined as the distance from neural lamella to the border of SPG with the underlying cells. We found that older foragers had a significantly higher mean depth blood-brain barrier than younger foragers. Depth measurements were made across 3 young foragers (n = 127) and 3 old foragers (n = 167). Data was log transformed prior to performing a Welch’s t-test (t(285.4) = 3.59, p<0.0005). Y-axis depicts depth range along a log transformed scale. Asterisk indicates significance.

## Discussion

This study presents a global anatomical description of the adult honeybee blood-brain barrier at the ultrastructural level. Additionally, we employed a systematic unbiased approach to compare ultrastructural features of the blood-brain barrier between a young forager (<15 days) group and an older forager group (≥15 days). Our analyses confirm that the cellular structure of the honeybee blood-brain barrier matches other described insect blood-brain barriers. Our use of a comprehensive, random, unbiased sampling regime led us to observe regions of the blood-brain barrier that might not be captured in more targeted sampling. Thus, we’ve revealed several structural details at the blood-brain barrier that warrant further study to understand full structural and functional diversity of the honeybee blood-brain barrier.

The honeybee blood-brain barrier metabolically and physically compartmentalizes the neuronal regions of the CNS from the hemolymph. It is essentially composed of the neural lamella and two surface glial subtypes, perineurial glia, and subperineurial glia. These components collectively regulate the import of nutrients and signals and the export of metabolic waste from the CNS as occurs in other insects (Maddrell and Treherne, 1967; Limmer *et al*., 2014). An aspect of the honeybee blood-brain barrier which was partially elucidated here but warrants further study is its heterogeneity. This heterogeneity mirrors, and may be directly related to, variation in the neuroglial composition of the regions directly underlying the surface glia layer.

Based on our morphological analysis, we confirmed that PG form the outermost layer of the blood-brain barrier in honeybees. These cells are positioned between the neural lamella and the SPG. Our understanding of PG function in honeybees is limited, and general knowledge about this cell type is derived from studies on *Drosophila* and few other insects. PG contribute to the formation of the neural lamella by depositing fibers that give it both stiffness and shape (Yildirim *et al*., 2019; Dunton *et al*., 2021). Their relatively small size and junctions oriented towards the hemolymph suggest that PG may act as an initial filter to prevent the entry of large molecules into the CNS interstitial space (Pogodalla, Winkler and Klämbt, 2022). They have direct contact with hemolymph that is filtered through the neural lamella and contain a conserved set of transporter proteins involved in nutrient uptake and efflux of toxins and other nonessential molecules (DeSalvo *et al*., 2014).

Although PG do not directly interact with neurons, they play a crucial role in neuronal development and survival. During embryogenesis, PG express a heparan sulfate proteoglycan that is vital for the proliferation of neuroblasts, suggesting a bidirectional communication system between PG and neuroblasts (Kanai *et al*., 2018). In the later stages of *Drosophila’s* lifespan, PG contribute to neuronal homeostasis and survival by taking up and transporting trehalose, the primary metabolic sugar substrate in insects (Volkenhoff *et al*., 2018). Their location and known functions implicate PG as a global hemolymph sensor, capable of perceiving and transducing information about the physiological state of the animal to other regions of the brain (Desalvo *et al*., 2011; Contreras and Sierralta, 2022). This ability to sense and communicate signals may contribute to the overall coordination of the central nervous system’s response to changes in the internal environment.

Despite their metabolic and signaling channels directly with hemolymph, PG are not tightly bound to one another, and thus unable to prevent small molecules from breaching their intracellular space. This essential diffusion barrier function is accomplished by the SPG, which form the layer immediately deep to the PG. SPG are large (up to 50 μm in the XY plane) polyploid cells with extensive protrusions that interdigitate with neighboring cells (Pogodalla, Winkler and Klämbt, 2022). This extensive interdigitation forms a tortuous maze of intercellular space, and greatly increases the diffusion distance that molecules would travel to enter the interstitial space. Studies in *Drosophila* have revealed that SPG form continuous segments of septate junctions that are crucial to the diffusion barrier between brain and hemolymph (Li *et al*., 2021). The structure and junctional contacts of SPG prevents paracellular diffusion, forcing the transport path of hydrophilic molecules to the array of transporters present in the SPG (DeSalvo *et al*., 2014; Limmer *et al*., 2014; Hindle *et al*., 2017).

To accommodate the trafficking of nutrients between the nervous system and circulatory system, the blood-brain barrier cells must produce both the molecular machinery and energy for diverse active transport mechanisms. Our results demonstrate that the honeybee blood-brain barrier has a relatively high volume fraction of both nuclei and mitochondria, values primarily driven by the SPG. The V_vmit_ of the honeybee blood-brain barrier is slightly less than that of V_vmit_ of honeybee flight muscle, one of the most metabolically active tissue types in the animal kingdom (Barth, 1992; Suarez *et al*., 2000; Fogarty *et al*., 2021). To our knowledge, this is the first direct analysis of the nuclear volume fraction of an insect blood-brain barrier, however the *Drosophila* blood-brain barrier is known to be both polyploid and multinucleate (Unhavaithaya and Orr-Weaver, 2012). We did not find a significant difference in either mitochondrial nor nuclear volume fraction between young and old foragers.

These subcellular characteristics likely exist to support the seemingly massive load of molecular transport that must happen between the CNS and hemolymph. Other than the import of nutritional molecules and the continuous maintenance of septate junctions, the SPG layer must facilitate the export of metabolic waste from the brain and host a suite of efflux transporters that capture and eject molecules back into circulation (DeSalvo *et al*., 2014). The energy output of SPG mitochondria likely supports an extensive repertoire of transport proteins and systems that facilitate the aforementioned barrier functions. Although there is lack of genomic information about honeybee SPG, transporters identified in transcriptomic analyses of *Drosophila* surface glial cells reveal a swath of evolutionary conserved proteins (DeSalvo *et al*., 2014). Further, a recent genomic analysis of halictid bees identified key nutrient transporters in the SPG which may be involved in organizing behaviors in highly social halictid species (Jones *et al*., 2023).

The majority of studies on the insect blood-brain barrier have implemented immunofluorescent imaging strategies and have focused on early life stage specimens. While these approaches have revealed numerous insights into the development and fundamental features and functions of surface glial cells, our randomized, unbiased ultrastructural imaging approach has captured the spatial heterogeneity of the adult honeybee blood-brain barrier. In doing so, our images reveal interesting blood-brain barrier structural details at certain locations around the brain. In an attempt to characterize the temporal heterogeneity of the adult honeybee blood-brain barrier, we compared specific ultrastructural features between young and old foragers, the latter of which are more likely to be undergoing senescence in the brain than the former.

The most heterogeneous structural feature we examined was the depth of the barrier, defined as the distance from the apical membrane of the PG to the basal membrane of the SPG. We observed great variance in barrier depth across all samples, with a minimum measured depth of 0.345 μm, to a maximum of 27.676 μm. Interestingly, we found that the mean blood-brain barrier depth of old foragers is larger than that of young foragers (Figure 6). An increase in cell number or cell volume could explain an age-related enlargement of blood-brain barrier depth. While neurogenesis does not occur in the adult honeybee brain, discrete neuropil regions of the forager brain increase in size with foraging and olfactory experience (Fahrbach, Strande and Robinson, 1995; Winnington, Napper and Mercer, 1996; Farris, Robinson and Fahrbach, 2001; Ismail, Robinson and Fahrbach, 2006).

It is possible that the blood-brain barrier also thickens with age. For example, the blood-brain barrier may increase water retention as a store for the brain as honeybee foragers lose hemolymph volume with age (Crailsheim, 1985). PG and/or SPG cells may also enlarge with foraging age as a protective measure against environmental toxins (Klein *et al*., 2017). While both are plausible, we suggest that the large average blood-brain barrier depth of old foragers may instead be a feature of naturally resilient foragers. The mortality rate of foragers increases with age, and thus our sampling method inherently contains a selection bias for old foragers capable of surviving ≥15 days of foraging (Dukas, 2008). Senescence in the honeybee brain manifests as increased oxidative stress-induced damage, downregulation of proteins involved in synaptic and intracellular signaling, and decreased learning and memory capabilities (Seehuus, Krekling and Amdam, 2006; Behrends *et al*., 2007; Scheiner and Amdam, 2009; Baker, Wolschin and Amdam, 2012). Age-related neural and cognitive dysregulation is further compounded by the stressors in a forager’s environment, such as their increased flight activity and exposure to environmental toxins (Tolfsen *et al*., 2011; Krupke *et al*., 2012; Williamson and Wright, 2013; Drummond *et al*., 2017). An increased blood-brain barrier diffusion distance could thus alleviate the added strain of hemolymph-derived factors which could increase the progression of brain senescence, thereby decreasing mortality risk.

To retain an unbiased tissue sampling approach, we did not track the regions of the brain in which stereological measurements were made. However, we would note observations from the extensive sampling and imaging required to set up this experiment that there is a marked difference in blood-brain barrier depth between the optic lobes and central regions of the brain. It is thus likely that the variance we measured in depth is linked to the fact that each of these regions of the brain had an equal chance of being measured. While the values we report do reflect that of the entire brain, it should be noted that the brain has two optic lobes, but one central region. Thus, for certain sectioning orientations, there may have been more blood-brain barrier samples derived from the optic lobes than the central body.

The honeybee optic lobes comprise three neuropils comprising axonal and dendritic tracts that process and transmit visual information from the retinas to the central regions, where it is integrated with other sensory information (Strausfeld, 2009). Here, the blood-brain barrier tended to be thick (∼10 μm), and dense with mitochondria and reticulated SPG membrane folds. The structural features may represent redundant paracellular and transcellular barrier mechanisms to maintain an optimal ionic environment for the transmission of visual information from the retinas to the synaptically dense central integratory regions. In contrast, the blood-brain barrier around central regions tended to be thin (∼1-5 μm). This structural form may enable the faster import of nutrients to accommodate the energetic demands of high synaptic activity, mirroring the increased capillary density in synaptically dense grey matter mammals (Wilhelm *et al*., 2016). These may reflect convergent strategies between the insect and mammalian blood-brain barrier to match the metabolic needs of the underlying nervous tissue (Villabona-Rueda *et al*., 2019). However, this structural form may also increase the susceptibility of these regions to harmful factors in hemolymph.

An interesting example of blood-brain barrier structural plasticity was described in a *Drosophila* septate junction mutant model (Babatz and Naffin, 2018). In this model, the unfolding of septate junctions is compromised as the brain grows in size during development. However, mutants survive to adulthood with few defects in blood-brain barrier integrity due to an increase in SPG interdigitation. Presumably, the lengthened diffusion pathway between the hemolymph and nervous tissue provided by the increased interdigitation compensates for mutation-induced loss in septate junctions (i.e. paracellular integrity). Interdigitation is an ancestral blood-brain barrier strategy to achieve a tight diffusion barrier around the brain (Bundgaard and Abbott, 2008). While this plasticity appears to be a compensatory mechanism to an experimental disruption, it is possible that honeybees employ both strategies to maintain integrity around different brain regions. It will be interesting for future analyses to determine if there is variation in the expression level of septate junction proteins and other integrity-associated proteins between the optic lobe and central bodies that could explain how barrier function is maintained with such variation in structure.

We made two notable observations within the regions of thin blood-brain barrier. First, we observed portions of the blood-brain barrier that are composed of a single protrusion from a surface glial cell (Figure 3A). We speculate that the identity of the cells that compose these thin regions are SPG cells based on several features. First, the morphology of the single-layer protrusions aligns with the characteristics of SPG. Secondly, the septate junctions that connect SPG cells are believed to create the tight diffusion barrier, indicating their likely involvement in this minimum requirement of the blood-brain barrier (Banerjee, Sousa and Bhat, 2006; Banerjee *et al*., 2008). Additionally, during embryo stages, SPG cells form the single surface glial layer, suggesting that this single-layer coverage may persist into adulthood in honeybees (Babatz, Naffin and Klämbt, 2018). Confirming this would be significant in understanding the distinct functions of PG and SPG in the blood-brain barrier and overall brain function, as well as elucidating the heterogeneity and division of labor among cell types in the insect blood-brain barrier.

We also observed at least one point in the blood-brain barrier where there appears to be a complete absence of surface glia. Figure 4C depicts a cellular region that lies deep to surface glia in the rest of the image but in one place breaches the surface glia to interface directly with the neural lamella. While the neural lamella provides some barrier function and protection, this point would be largely exposed to hemolymph *in vivo*. There are multiple regions of mammalian blood-brain barrier which allow nearly free exchange between the blood and CNS, enabling the sensing of various factors in the blood (Wilhelm *et al*., 2016). Regions of the insect CNS without a glial blood-brain barrier may act in a similar fashion, with the neural lamella still providing a degree of protection. Despite the seemingly weak blood-brain barrier protection that these regions may have, we posit that there are enough protection systems in place to at least maintain the proper ionic milieu for electrochemical signaling. This is the most essential blood-brain barrier function across species with a centralized nervous system, and the loss of this function would render the tissue inoperable. The potential absence of PG in regions with a single layer blood-brain barrier suggest their function to the operation of the barrier is not essential, but also that they play a significant role in regions where they are present. Their presence seems to be most pronounced at points where cortex-enwrapped neuronal cell bodies are present beneath the blood-brain barrier and further studies are warranted to understand their ability to sense and propagate signals from the circulatory system towards these and other integration centers in the brain.

## References

Abbott, N. J. (1992) ‘Comparative Physiology of the Blood-Brain Barrier’, in Advances in Experimental Medicine and Biology, pp. 371–396. doi: 10.1007/978-3-642-76894-1_15.

Abbott, N. J. et al. (2010) ‘Structure and function of the blood–brain barrier’, Neurobiology of Disease. Elsevier Inc., 37(1), pp. 13–25. doi: 10.1016/j.nbd.2009.07.030.

Abbott, N. J., Lane, N. J. and Bundgaard, M. (1986) ‘The Blood-Brain Interface in Invertebrates’, Ann. N. Y. Acad. Sci., 481, pp. 20–42.

Andersson, O. et al. (2013) ‘The Grasshopper: A Novel Model for Assessing Vertebrate Brain Uptake’, Journal of Pharmacology and Experimental Therapeutics, 346(2), pp. 211–218. doi: 10.1124/jpet.113.205476.

Avarguès-Weber, A. and Giurfa, M. (2013) ‘Conceptual learning by miniature brains’, Proceedings of the Royal Society B: Biological Sciences, 280(1772), pp. 19–21. doi: 10.1098/rspb.2013.1907.

Awasaki, T. et al. (2008) ‘Organization and Postembryonic Development of Glial Cells in the Adult Central Brain of Drosophila’, The Journal of Neuroscience, 28(51), pp. 13742– 13753. doi: 10.1523/JNEUROSCI.4844-08.2008.

Babatz, F. and Naffin, E. (2018) ‘The Drosophila Blood-Brain Barrier Adapts to Cell Growth by Unfolding of Pre-existing Septate Junctions’, pp. 1–14. doi: 10.1016/j.devcel.2018.10.002.

Babatz, F., Naffin, E. and Klämbt, C. (2018) ‘The Drosophila Blood-Brain Barrier Adapts to Cell Growth by Unfolding of Pre-existing Septate Junctions’, Developmental Cell, 47(6), pp. 697–710.e3. doi: 10.1016/j.devcel.2018.10.002.

Baker, N., Wolschin, F. and Amdam, G. V. (2012) ‘Age-related learning deficits can be reversible in honeybees Apis mellifera’, Experimental Gerontology. Elsevier Inc., 47(10), pp. 764–772. doi: 10.1016/j.exger.2012.05.011.

Banerjee, S. et al. (2008) ‘Septate junctions are required for ommatidial integrity and blood-eye barrier function in Drosophila’, Developmental Biology, 317(2), pp. 585–599. doi: 10.1016/j.ydbio.2008.03.007.

Banerjee, S., Sousa, A. D. and Bhat, M. A. (2006) ‘Organization and function of septate junctions: An evolutionary perspective’, Cell Biochemistry and Biophysics, 46(1), pp. 65–77. doi: 10.1385/CBB:46:1:65.

Barth, E. (1992) ‘Ultrastructural quantitation of mitochondria and myofilaments in cardiac muscle from 10 different animal species including man’, Journal of Molecular and Cellular Cardiology, 24(7), pp. 669–681. doi: 10.1016/0022-2828(92)93381-S.

Behrends, A. et al. (2007) ‘Cognitive aging is linked to social role in honey bees (Apis mellifera)’, 42, pp. 1146–1153. doi: 10.1016/j.exger.2007.09.003.

Bundgaard, M. and Abbott, N. J. (2008) ‘All vertebrates started out with a glial blood-brain barrier 4-500 million years ago’, Glia, 56(7), pp. 699–708. doi: 10.1002/glia.20642.

Carlson, S. D. et al. (2000) ‘Blood Barriers of the Insect’, Annual Review of Entomology, 45(1), pp. 151–174. doi: 10.1146/annurev.ento.45.1.151.

Chen, M. et al. (2019) ‘Brain Endothelial Cells are Exquisite Sensors of Age-Related Circulatory Cues’, SSRN Electronic Journal. doi: 10.2139/ssrn.3406390.

Contreras, E. G. and Klämbt, C. (2023) ‘The Drosophila blood-brain barrier emerges as a model for understanding human brain diseases’, Neurobiology of Disease, 180(December 2022), p. 106071. doi: 10.1016/j.nbd.2023.106071.

Contreras, E. G. and Sierralta, J. (2022) ‘The Fly Blood-Brain Barrier Fights Against Nutritional Stress’, Neuroscience Insights, 17, p. 263310552211202. doi: 10.1177/26331055221120252.

Cook, C. N. et al. (2020) ‘Individual learning phenotypes drive collective behavior’, Proceedings of the National Academy of Sciences of the United States of America, 117(30), pp. 17949–17956. doi: 10.1073/pnas.1920554117.

Crailsheim, K. (1985) ‘Distribution of haemolymph in the honeybee (Apis mellifica) in relation to season, age and temperature’, Journal of Insect Physiology, 31(9), pp. 707– 713. doi: 10.1016/0022-1910(85)90051-4.

Desalvo, M. K. et al. (2011) ‘Physiologic and anatomic characterization of the brain surface glia barrier of Drosophila’, Glia, 59(9), pp. 1322–1340. doi: 10.1002/glia.21147.

DeSalvo, M. K. et al. (2014) ‘The Drosophila surface glia transcriptome: evolutionarily conserved blood-brain barrier processes’. Available at: https://www.researchgate.net/profile/Zeid_Rusan/publication/268875882_The_Drosophila_surface_glia_transcriptome_Evolutionary_conserved_blood-brain_barrier_processes/links/54ad91e80cf2213c5fe412b6.pdf (Accessed: 19 April 2016).

Drummond, J. et al. (2017) ‘Spontaneous honeybee behaviour is altered by persistent organic pollutants’, Ecotoxicology. Springer US, pp. 141–150. doi: 10.1007/s10646-016-1749-0.

Dukas, R. (2008) ‘Mortality rates of honey bees in the wild’, Insectes Sociaux, 55(3), pp. 252–255. doi: 10.1007/s00040-008-0995-4.

Dunton, A. D. et al. (2021) ‘Form and Function of the Vertebrate and Invertebrate Blood-Brain Barriers’, International Journal of Molecular Sciences, 22(22), p. 12111. doi: 10.3390/ijms222212111.

Ehrlich, P. (1885) *Das sauerstufbudurfnis des organismus,, Eine Farbenanalytische Studie*. Berlin: Hirschwald.

Fahrbach, S. E., Strande, J. L. and Robinson, G. E. (1995) ‘Neurogenesis is absent in the brains of adult honey bees and does not explain behavioral neuroplasticity’, 197, pp. 145–148.

Farris, S. M., Robinson, G. E. and Fahrbach, S. E. (2001) ‘Experience- and Age-Related Outgrowth of Intrinsic Neurons in the Mushroom Bodies of the Adult Worker

Honeybee’, The Journal of Neuroscience, 21(16), pp. 6395–6404. doi: 10.1523/JNEUROSCI.21-16-06395.2001.

Fogarty, M. J. et al. (2021) ‘Quantifying mitochondrial volume density in phrenic motor neurons’, Journal of Neuroscience Methods. Elsevier B.V., 353(January), p. 109093. doi: 10.1016/j.jneumeth.2021.109093.

Freeman, M. R. (2015) ‘Drosophila central nervous system glia’, Cold Spring Harbor Perspectives in Biology, 7(11). doi: 10.1101/cshperspect.a020552.

Goldmann, E. (1913) ‘Vitalfärbung am Zentralnervensystem : Beitrag zur Physio-Pathologie des Plexux Chorioideus und der Hirnhäute.’

Griffith, J. I. et al. (2020) ‘Addressing BBB heterogeneity: A new paradigm for drug delivery to brain tumors’, Pharmaceutics, 12(12), pp. 1–38. doi: 10.3390/pharmaceutics12121205.

Groh, C. and Rössler, W. (2020) ‘Analysis of Synaptic Microcircuits in the Mushroom Bodies of the Honeybee’, Insects, 11(1), p. 43. doi: 10.3390/insects11010043.

Habenstein, J. et al. (2023) ‘3D atlas of cerebral neuropils with previously unknown demarcations in the honey bee brain’, Journal of Comparative Neurology, 531(11), pp. 1163–1183. doi: 10.1002/cne.25486.

Hartung, D. A. K., Kirkton, S. D. and Harrison, J. F. (2004) ‘Ontogeny of tracheal system structure: A light and electron-microscopy study of the metathoracic femur of the American locust, Schistocerca americana’, Journal of Morphology, 262(3), pp. 800–812. doi: 10.1002/jmor.10281.

Hindle, S. J. et al. (2017) ‘Evolutionarily Conserved Roles for Blood-Brain Barrier Xenobiotic Transporters in Endogenous Steroid Partitioning and Behavior’, Cell Reports. ElsevierCompany., 21(5), pp. 1304–1316. doi: 10.1016/j.celrep.2017.10.026.

Hurst, E. W. and Davies, O. L. (1950) ‘Studies on the Blood-Brain-Barrier. II. Attempts to Influence the Passage of Substances into the Brain’, British Journal of Pharmacology and Chemotherapy, 5(2), pp. 147–164. doi: 10.1111/j.1476-5381.1950.tb01002.x.

Ismail, N., Robinson, G. E. and Fahrbach, S. E. (2006) ‘Stimulation of muscarinic receptors mimics experience-dependent plasticity in the honey bee brain’, Proceedings of the National Academy of Sciences, 103(1), pp. 207–211. doi: 10.1073/pnas.0508318102.

Ito, K. et al. (2014) ‘A Systematic Nomenclature for the Insect Brain’, Neuron, 81(4), pp. 755–765. doi: 10.1016/j.neuron.2013.12.017.

Jones, B. M. et al. (2023) ‘Convergent and complementary selection shaped gains and losses of eusociality in sweat bees’, Nature Ecology & Evolution. Springer US, 7(4), pp. 557–569. doi: 10.1038/s41559-023-02001-3.

Kaiser, A. et al. (2022) ‘A three-dimensional atlas of the honeybee central complex, associated neuropils and peptidergic layers of the central body’, Journal of Comparative Neurology, 530(14), pp. 2416–2438. doi: 10.1002/cne.25339.

Kanai, M. I. et al. (2018) ‘Regulation of neuroblast proliferation by surface glia in the Drosophila larval brain’, Scientific Reports, 8(1), p. 3730. doi: 10.1038/s41598-018-22028-y.

Kenyon, F. (1896) ‘The brain of the bee. A preliminary contribution to the morphology of the nervous system of the arthropoda.’, J Comp Neurol, 6, pp. 133–210.

Klein, S. et al. (2017) ‘Why Bees Are So Vulnerable to Environmental Stressors’, Trends in Ecology & Evolution. Elsevier Ltd, 32(4), pp. 268–278. doi: 10.1016/j.tree.2016.12.009.

Kraft, N. et al. (2023) ‘Expansion microscopy in honeybee brains for high-resolution neuroanatomical analyses in social insects’, Cell and Tissue Research. Springer Berlin Heidelberg, (0123456789). doi: 10.1007/s00441-023-03803-4.

Kremer, M. C. et al. (2017) ‘The glia of the adult Drosophila nervous system’, GLIA, 65(4), pp. 606–638. doi: 10.1002/glia.23115.

Krupke, C. H. et al. (2012) ‘Multiple routes of pesticide exposure for honey bees living near agricultural fields’, PLoS ONE, 7(1). doi: 10.1371/journal.pone.0029268.

Lewandowsky, M. (1900) ‘Zur lehre von der cerebrospinalflussigkeit’, Z Klin Med, 40, pp. 480–494.

Li, X. et al. (2021) ‘The cAMP effector PKA mediates Moody GPCR signaling in Drosophila blood-brain barrier formation and maturation’, bioRxiv.

Limmer, S. et al. (2014) ‘The Drosophila blood-brain barrier: development and function of a glial endothelium.’, Frontiers in neuroscience, 8, p. 365. doi: 10.3389/fnins.2014.00365.

Maddrell, S. H. P. and Treherne, J. E. (1967) ‘The Ultrastructure of the Perineurium in two Insect Species, Carausius Morosus and Periplaneta Americana’, Journal of Cell Science, 2(1), pp. 119–128. doi: 10.1242/jcs.2.1.119.

Marcos, R., Monteiro, R. A. F. and Rocha, E. (2012) ‘The use of design-based stereology to evaluate volumes and numbers in the liver: a review with practical guidelines’, Journal of Anatomy, 220(4), pp. 303–317. doi: 10.1111/j.1469-7580.2012.01475.x.

Mayhew, T. M. and Orive, L. -M. C. (1974) ‘Caveat on the use of the Delesse principle of areal analysis for estimating component volume densities’, Journal of Microscopy, 102(2), pp. 195–207. doi: 10.1111/j.1365-2818.1974.tb03979.x.

Menzel, R. (2001) ‘Searching for the memory trace in a mini-brain, the honeybee.’, *Learning & memory (Cold Spring Harbor*, N.Y.), 8(2), pp. 53–62. doi: 10.1101/lm.38801.

Menzel, R. (2012) ‘The honeybee as a model for understanding the basis of cognition’, Nature Reviews Neuroscience. Nature Publishing Group, 13(11), pp. 758–768. doi: 10.1038/nrn3357.

Menzel, R. and Giurfa, M. (2001) ‘Cognitive architecture of a mini-brain: The honeybee’, Trends in Cognitive Sciences, 5(2), pp. 62–71. doi: 10.1016/S1364-6613(00)01601-6.

Mironov, A. (2017) ‘Stereological morphometric grids for ImageJ’, Ultrastructural Pathology, 41(1), pp. 126–126. doi: 10.1080/01913123.2016.1272665.

Munch, D. et al. (2013) ‘Obtaining Specimens with Slowed, Accelerated and Reversed Aging in the Honey Bee Model’, Journal of visualized experiments, (78), p. e50550. doi: doi:10.3791/50550.

Noumbissi, M. E., Galasso, B. and Stins, M. F. (2018) ‘Brain vascular heterogeneity: implications for disease pathogenesis and design of in vitro blood–brain barrier models’, Fluids and Barriers of the CNS. BioMed Central, 15(1), p. 12. doi: 10.1186/s12987-018-0097-2.

Paoli, M. and Galizia, G. C. (2021) ‘Olfactory coding in honeybees’, Cell and Tissue Research. Springer Berlin Heidelberg, 383(1), pp. 35–58. doi: 10.1007/s00441-020-03385-5.

Pivoriūnas, A. and Verkhratsky, A. (2021) ‘Astrocyte–Endotheliocyte Axis in the Regulation of the Blood–Brain Barrier’, Neurochemical Research. Springer US, 46(10), pp. 2538–2550. doi: 10.1007/s11064-021-03338-6.

Pogodalla, N., Winkler, B. and Klämbt, C. (2022) ‘Glial Tiling in the Insect Nervous System’, Frontiers in Cellular Neuroscience, 16(February), pp. 1–7. doi: 10.3389/fncel.2022.825695.

Quigley, T. P., Amdam, G. V. and Rueppell, O. (2018) ‘Honeybee Workers as Models of Aging’, in Conn’s Handbook of Models for Human Aging. Second Edi. Elsevier, pp. 533–547. doi: 10.1016/B978-0-12-811353-0.00040-3.

Ribatti, D. et al. (2006) ‘Development of the blood-brain barrier: A historical point of view’, Anatomical Record - Part B New Anatomist, 289(1), pp. 3–8. doi: 10.1002/ar.b.20087.

Rybak, J. (2012) ‘The Digital Honey Bee Brain Atlas’, in Galizia, C. G., Eisenhardt, D., and Giurfa, M. (eds) Honeybee Neurobiology and Behavior. Dordrecht: Springer Netherlands, pp. 125–140. doi: 10.1007/978-94-007-2099-2_11.

Santuy, A. et al. (2018) ‘A quantitative study on the distribution of mitochondria in the neuropil of the juvenile rat somatosensory cortex’, Cerebral Cortex, 28(10), pp. 3673– 3684. doi: 10.1093/cercor/bhy159.

Saunders, N. R. et al. (2014) ‘The rights and wrongs of blood-brain barrier permeability studies: a walk through 100 years of history’, Frontiers in Neuroscience, 8(DEC), pp. 1–26. doi: 10.3389/fnins.2014.00404.

Scheiner, R. et al. (2013) ‘Standard methods for behavioural studies of Apis mellifera’, Journal of Apicultural Research, 52(4), pp. 1–58. doi: 10.3896/IBRA.1.52.4.04.

Scheiner, R. and Amdam, G. V. (2009) ‘Impaired tactile learning is related to social role in honeybees.’, The Journal of experimental biology, 212(Pt 7), pp. 994–1002. doi: 10.1242/jeb.021188.

Schofield, P. K. and Treherne, J. E. (1984) ‘Localization of the blood-brain barrier of an insect: Electrical model and analysis’, The Journal of experimental biology, 109, pp. 319–331.

Seehuus, S. C., Krekling, T. and Amdam, G. V. (2006) ‘Cellular senescence in honey bee brain is largely independent of chronological age’, Experimental Gerontology, 41(11), pp. 1117–1125. doi: 10.1016/j.exger.2006.08.004.

Seeley, T. D. (1986) ‘Social foraging by honey bees: How colonies allocate foraging among patches of flowers.’, Behavioral Ecology and Sociobiology, 19(5), pp. 343–354.

Siefert, P., Buling, N. and Grünewald, B. (2021) ‘Honey bee behaviours within the hive: Insights from long-term video analysis’, PLoS ONE, 16(3 March), pp. 1–14. doi: 10.1371/journal.pone.0247323.

Sommerlandt, F. M. J. et al. (2019) ‘Immediate early genes in social insects: a tool to identify brain regions involved in complex behaviors and molecular processes underlying neuroplasticity’, Cellular and Molecular Life Sciences. Springer International Publishing, pp. 637–651. doi: 10.1007/s00018-018-2948-z.

Stern, L. (1934) ‘A propos de la méthod d’investigation du fonctionnement de la barrière hémato-encéphalique’, C R Soc Biol, 115, pp. 1059–1061.

Stern, L. and Gautier, R. (1918) ‘Le passage dans le liquide céphalo-rachidien de substances introduites dans la circulation et leur action sur le système nerveux central chez les différentes espèces animales’, R C R d Ia Soc Phys d’hist nat Genève, 35, pp. 91–94.

Strausfeld, N. J. (2009) ‘Brain and Optic Lobes’, in *Encyclopedia of Insects*. Second Edi. Elsevier, pp. 121–130. doi: 10.1016/B978-0-12-374144-8.00042-4.

Suarez, R. K. et al. (2000) ‘Mitochondrial function in flying honeybees (Apis mellifera): Respiratory chain enzymes and electron flow from complex III to oxygen’, Journal of Experimental Biology, 203(5), pp. 905–911.

Tolfsen, C. C. et al. (2011) ‘Flight restriction prevents associative learning deficits but not changes in brain protein-adduct formation during honeybee ageing.’, The Journal of experimental biology, 214, pp. 1322–32. doi: 10.1242/jeb.049155.

Unhavaithaya, Y. and Orr-Weaver, T. L. (2012) ‘Polyploidization of glia in neural development links tissue growth to blood – brain barrier integrity’, Genes & Development, pp. 31–36. doi: 10.1101/gad.177436.111.Freely.

Vein, A. A. (2008) ‘Science and Fate: Lina Stern (1878–1968), A Neurophysiologist and Biochemist’, Journal of the History of the Neurosciences, 17(2), pp. 195–206. doi: 10.1080/09647040601138478.

Villabona-Rueda, A. et al. (2019) ‘The Evolving Concept of the Blood Brain Barrier (BBB): From a Single Static Barrier to a Heterogeneous and Dynamic Relay Center’, Frontiers in Cellular Neuroscience, 13(September). doi: 10.3389/fncel.2019.00405.

Volkenhoff, A. et al. (2018) ‘Live imaging using a FRET glucose sensor reveals glucose delivery to all cell types in the Drosophila brain’, Journal of Insect Physiology. Elsevier, 106(February), pp. 55–64. doi: 10.1016/j.jinsphys.2017.07.010.

Weiler, A. et al. (2017) ‘Metabolite transport across the mammalian and insect brain diffusion barriers’, Neurobiology of Disease. Elsevier Inc., 107, pp. 15–31. doi: 10.1016/j.nbd.2017.02.008.

Wilhelm, I. et al. (2016) ‘Heterogeneity of the blood-brain barrier’, Tissue Barriers, 4(1), p. e1143544. doi: 10.1080/21688370.2016.1143544.

Williamson, S. M. and Wright, G. A. (2013) ‘Exposure to multiple cholinergic pesticides impairs olfactory learning and memory in honeybees’, pp. 1799–1807. doi: 10.1242/jeb.083931.

Winnington, A. P., Napper, R. M. and Mercer, A. R. (1996) ‘Structural plasticity of identified glomeruli in the antennal lobes of the adult worker honey bee’, The Journal of Comparative Neurology, 365(3), pp. 479–490. doi: 10.1002/(SICI)1096-9861(19960212)365:3<479::AID-CNE10>3.0.CO;2-M.

Winston, M. L. (1987) The Biology of the Honey Bee. Cambridge, MA: Harvard University Press.

Wolschin, F., Münch, D. and Amdam, G. V. (2009) ‘Structural and proteomic analyses reveal regional brain differences during honeybee aging’, Journal of Experimental Biology, 212(24), pp. 4027–4032. doi: 10.1242/jeb.033845.

Yildirim, K. et al. (2019) ‘Drosophila glia: Few cell types and many conserved functions’, GLIA, pp. 5–26. doi: 10.1002/glia.23459.

